# Profiling of presynaptic mRNAs reveals a role for axonal PUMILIOs in associative memory formation

**DOI:** 10.1101/733428

**Authors:** Rachel Arey, Rachel Kaletsky, Coleen T. Murphy

## Abstract

Presynaptic protein synthesis is important in the adult central nervous system; however, the set of mRNAs localized to presynaptic areas has yet to be identified in any organism. We differentially labeled somatic and synaptic compartments nervous system-wide in adult *C. elegans* and isolated synaptic regions for deep sequencing. Analysis of the synaptic transcriptome reveals that synaptic transcripts are predicted to have specialized functions in neurons. Differential expression analysis identified 543 genes enriched in synaptic regions relative to somatic regions, with synaptic functions conserved in higher organisms. We find that mRNAs for *pumilio* RNA-binding proteins are abundant in synaptic regions, which we confirmed through high-sensitivity in situ hybridization. We identified a new role for the PUM2 orthologs *puf-7/8* as repressors of memory formation through regulation of the mitochondrial dynamics regulator, *mff-1.* Identification of presynaptic mRNAs provides insight into mechanisms that regulate synaptic function and behavior.

## Introduction

Neurons are polarized, structurally complex cells comprised of functionally distinct compartments, with dendrites, somatic regions, and axon terminals often operating in different microenvironments (Glock et al., 2017). These compartments must often rapidly respond to and integrate discrete, spatially restricted stimuli. One of the mechanisms by which synapses coordinate dynamic responses is through localized protein synthesis in dendrites, as protein transport alone from the soma is too slow to meet timing demands of synaptic signaling. It was long thought that local translation in the adult brain exclusively occurred in postsynaptic, not presynaptic, compartments, due to the failure of electron microscopy studies to visualize polysomes in presynaptic terminals (Twiss and Fainzilber, 2009), though recent expansion microscopy techniques have found that ribosomes are indeed present in presynaptic terminals (Hafner et al., 2019).

The importance of dendritic local translation in neuronal plasticity was first identified when it was observed that protein-synthesis-dependent long-term potentiation (LTP) and long-term depression (LTD) occur when the soma is physically disconnected from postsynaptic regions (Cracco et al., 2005; Huber et al., 2000; Kang and Schuman, 1996). Dendritic local translation is also thought to play an important role in memory storage, as it provides a potential mechanism for the strengthening of specific synapses. There have been many recent advances in characterizing mRNAs localized to dendritic regions, including the identification of over 2000 synaptic mRNAs localized to the neuropil of the hippocampus (Cajigas et al., 2012) as well as compartment specific 3’ UTR usage (Tushev et al., 2018), and the development of new tools to study and visualize the translation and localization of specific mRNAs (reviewed in (Glock et al., 2017)).

More recently, localized protein synthesis has been revealed in axons and presynaptic terminals (Hafner et al., 2019; Jung et al., 2012). Specifically, local protein synthesis is important for the response of the axonal growth cone to guidance cues (Jung et al., 2012), and axonal translation of the t-SNARE protein SNAP25 was found to be necessary for the proper assembly of presynaptic terminals during development (Batista et al., 2017). In adulthood, axonal protein synthesis plays a critical role in response to nerve injury (Terenzio et al., 2018); mTOR is rapidly translated upon injury and regulates its own translation, as well as the levels of retrograde signaling proteins (Terenzio et al., 2018). The role of presynaptic protein synthesis in plasticity and behavior is less well characterized, though it is necessary for branch-specific long-term facilitation in *Aplysia* (Martin et al., 1997; Wang et al., 2009). More recent studies in the mammalian brain have found that long-term plasticity of GABA release (Younts et al., 2016) and neurotransmitter release at the calyx of Held (Scarnati et al., 2018) both involve presynaptic translation.

In order to further our understanding of how presynaptic protein synthesis regulates plasticity and behavior, it is critical to identify presynaptically localized transcripts. Recent studies identified the axonal transcriptome and translatome of cultured motor neurons (Nijssen et al., 2018) and retinal ganglion cells (Shigeoka et al., 2016), respectively; however, the full set of transcripts in the nervous system that are localized specifically to presynaptic compartments have yet to be described in any system. Furthermore, it is unknown if presynaptically localized transcripts contribute to complex behaviors.

We recently developed a technique to isolate and RNA-sequence specific tissues and neuronal subtypes in the nematode worm *C. elegans* (Kaletsky et al., 2016), revealing new regulators of neuron-specific phenotypes, such as axon regeneration and associative learning and memory. Here we describe how we have differentially labeled somatic, axonal, and presynaptic compartments of the adult *C. elegans* nervous system using a dual-fluorescent protein strategy. These differentially labeled compartments can be isolated by fluorescence activated cell sorting (FACS) and used to identify presynaptically-localized RNAs. We find that these genes have predicted synaptic functions, and their mammalian orthologs are known to function in synaptic and axonal regions, suggesting that the localization of the novel synaptic mRNAs identified here is likely conserved. We also find that synapse-enriched genes are predicted to bind mRNA, including a number of *Pumilio* orthologs (*pufs*) that regulate memory formation. Because the regulation of synaptic transmission and presynaptic function is highly conserved between *C. elegans* and mammals, these synaptically-localized transcripts likely function in higher organisms to regulate processes where local protein synthesis is required, such as repair and plasticity.

## Results

### Isolation of Synaptic transcripts using a dual-fluorescent labeling strategy

We previously developed a technique to identify the transcriptomes of adult *C. elegans* tissues (Kaletsky et al., 2016; Kaletsky et al., 2018) that utilizes tissue-specific labeling by promoter-driven fluorescent proteins, outer cuticle breaking, size-specific filtering and sorting of cells by FACS, which can used for RNA-isolation and transcriptome analysis by RNA-seq. Using this approach, we identified transcripts expressed in specific tissues and individual cell types and neurons (Kaletsky et al., 2016). We discovered that the enzymatic and mechanical dissociation steps of isolating neurons using this technique could result in their fragmentation; fluorescently labeled neurites were often observed during sample preparation (Figure 1A). Because neurites and synapses contain their own mRNA, we devised a strategy to take advantage of this fragmentation and isolate specific neuronal sub-compartments, differentially labeling neurons with different fluorescent protein markers, which enabled us to simultaneously collect somatic and presynaptic regions from the same adult neuronal populations for transcriptomic analysis. Somatic regions were labeled with mCherry under the control of the promoter for the pan-neuronal Rab family GTPase *rab-3*, while a RAB-3∷GFP translational fusion, which localizes to presynaptic regions and is a widely used synaptic marker (Mahoney et al., 2006), was used to specifically label synapses (Figure 1B). Differentially-localized fluorescent proteins were detectable by microscopy (Figure 1C), and upon performing flow cytometry of isolated neurons from *pRab-3∷mCherry;pRab-3∷RAB-3∷GFP* animals, GFP+ pre-synaptic regions were isolated independently of mCherry+ cell bodies, as well as double-positive (GFP+/mCherry+) events which likely contain intact axons (Figure 1D). Each of these three isolated populations contained RNA that could be isolated and subjected to RNA-seq. We generated six biological replicates of these differentially-labeled fluorescent samples. Principle components analysis (PCA) revealed that all six mCherry+ soma samples clustered together, while two of the six GFP+ synaptic samples appeared to be outliers and were discarded from further analysis (Figure S1A). The remaining samples clustered well by isolated subcompartment (Figure S1B).

**Figure 1.**
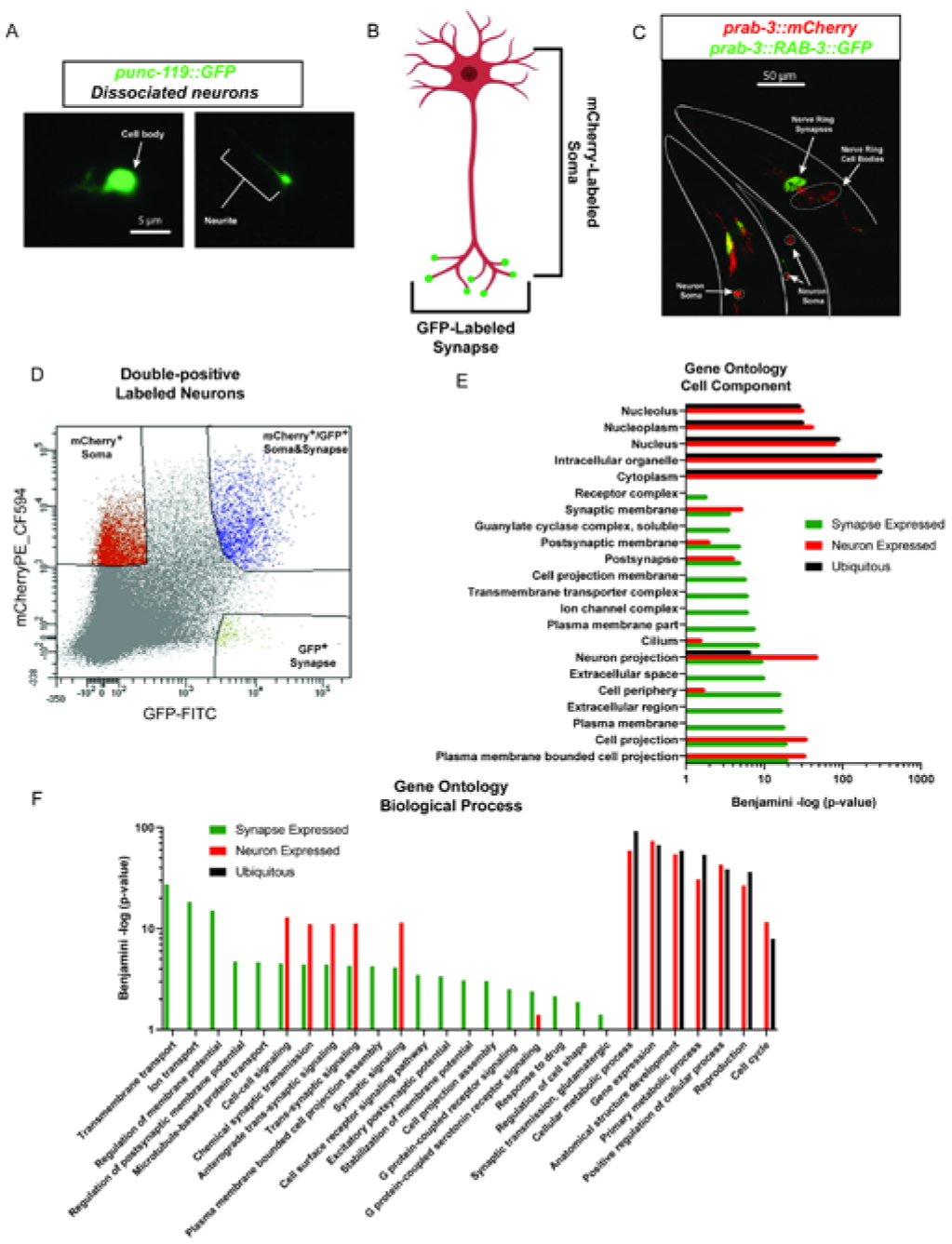
Isolation and RNA-seq of presynaptic mRNAs reveals specialized neuronal functions. **A)** Neuronal cell isolation results in fragmentation of cells, where cell bodies (left) can be detected separately from neurites (right). **B)** Schematic of the dual-fluorescent protein strategy for isolating synaptic and somatic compartments in *C. elegans.* **C)** Representative confocal images of *rab-3p∷mCherry; rab-3p∷RAB-3∷GFP* transgenic worms. Neurons in the head (outlined in white) and tail (outlined in gray) show distinct expression of fluorescent proteins. Cell bodies (outlined in white, arrows) express somatic mCherry (red), while nerve ring synapses exclusively express GFP (green, arrows. Colocalization of the two fluorescent proteins (yellow) is also evident. **D)** FACS plot displaying ability to isolate synaptic (GFP+, green), somatic (mCherry+, red), and synaptic+somatic (GFP+/mCherry+, blue) compartments using adult neuron cell isolation method (Kaletsky et al., 2016). **E-F)** Significant GO terms (padj<0.05) highlight that synapse-expressed genes(green) are predicted to be specialized neuronal genes in terms of both localization (**E**), enriched for membrane related terms and depleted for nuclear terms, unlike ubiquitous (black) and neuron-expressed genes (red) and function (**F**).

### Presynaptically expressed genes characterize synaptic function

In order to classify a gene as “expressed in synaptic compartments,” it had to have an average of 10 counts across the 4 remaining GFP+ synaptic samples. We additionally filtered out previously-identified ubiquitously-expressed genes, that are detected across all adult tissue samples (Kaletsky et al., 2018). Using these cutoffs, we identified 8,778 “synapse expressed” genes (Table S1). Comparison of GO terms of “synapse-expressed” genes (Table S1) to previously published neuron-expressed genes (Kaletsky et al., 2018) and ubiquitous genes (Kaletsky et al., 2018) revealed that “synapse-expressed” genes are predicted to have specialized neuronal functions that are synaptic in nature (Figure 1E,F), suggesting that the enrichment was successful.

For example, cellular component GO terms shared between neuron-enriched and synapse-expressed gene sets included *neuron projection* and *synaptic membrane*, but some terms were exclusively synaptic, such as *receptor complex, ion channel complex*, and *plasma membrane* (Figure 1E). Synapse-expressed genes were not predicted to function in other cellular components, such as the nucleus, which were highly significant in the neuron-expressed and ubiquitous gene lists (Figure 1E). Predicted functions exclusive to synapse-expressed genes included *ion transport, regulation of membrane potential, microtubule based protein transport*, and *synaptic transmission, glutamatergic* (Figure 1F). Shared terms with neuron-expressed genes included *synaptic signaling, chemical synaptic transmission*, and *cell-cell signaling*, but terms that denote nuclear functions such as *gene expression* are absent in synapse-expressed genes (Figure 1F). These results suggest that we have identified genes that contribute to well-characterized aspects of synaptic function

### Identification of Synaptic Differentially Expressed Genes (DEGs)

In addition to identifying mRNAs that were present at the synapse, we were particularly interested in finding mRNAs that were most significantly enriched in presynaptic regions. Therefore, we used DESeq2 for differential expression analysis (Love et al., 2014) to determine which transcripts were expressed at significantly higher levels (FDR <0.05) in synaptic samples (GFP^+^, Figure 1D) relative to somatic samples (mCherry^+^, Figure 1D), revealing 542 synaptic DEGs (Figure 2A, Table S2). These synaptic DEGs were significantly enriched (∼ 91%, p = 6.23 × 10^−60^, hypergeometric test) for previously-identified adult neuronal genes ((Kaletsky et al., 2016), Figure 2B), indicating that we had identified a subset of neuronal genes that are enriched in presynaptic regions.

**Figure 2.**
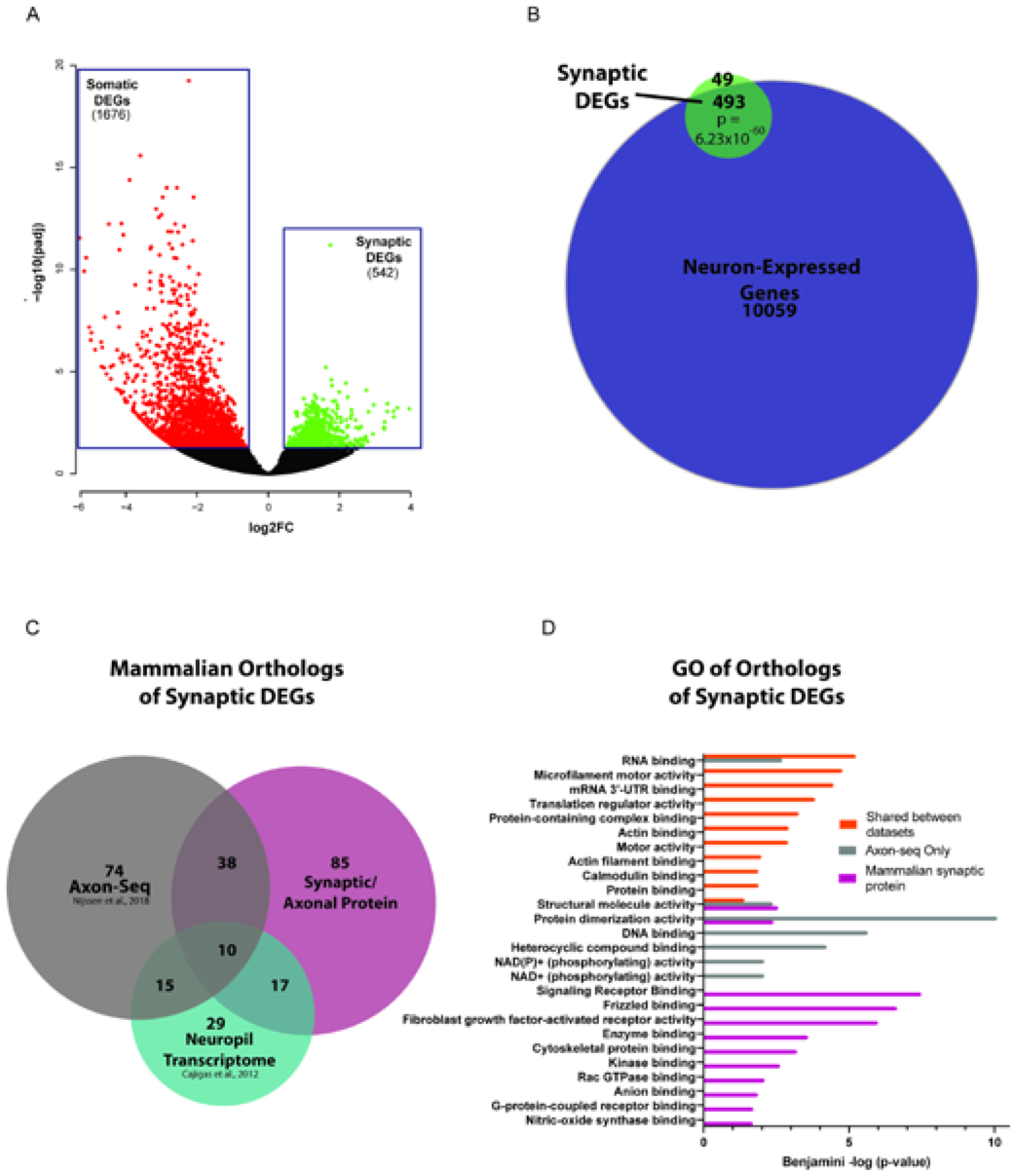
Characterization of Synaptic DEGs. **A)** Volcano plot of Synaptic DEGs (Green) relative to Somatic (Red) samples. FDR for DEGs = 0.05 **B)** Synaptic DEGs significantly overlap with previously identified adult-neuron expressed genes (Kaletsky et al., 2016) P-values: hypergeometric distributions. **C)** Mammalian orthologs of *C. elegans* synaptic DEGs function in synapses (for citations, see Table S3) or have been detected in previous synaptic neuropil ((Cajigas et al., 2012), blue) and axonal transcriptomic ((Nijssen et al., 2018), grey) datasets. **D)** GO analysis of known conserved synaptic DEGs reveals synaptic and axonal functions (orange, grey, magenta), and an enrichment of RNA binding proteins and translational regulators (orange).

### Synaptic DEGS have known synaptic and axonal functions in mammals

*C. elegans’* nervous system has a high degree of conservation with mammalian neurons. Using the OrthoList2 tool (Kim et al., 2018), 311 of the *C. elegans* synaptic DEGs have predicted mammalian orthologs (Table S3). Of those mammalian orthologs, 269 have been previously validated as axonal or synaptic: protein products of 150 genes were found either function in synapses and axons (Synaptic/Axonal Protein, Figure 2C), while previous transcriptomic studies that captured subsets of axonal or synaptic regions identified orthologs of 137 and 71 synaptic DEGS, respectively (Axon-seq and Neuropil transcriptome, Figure 2C, (Cajigas et al., 2012; Nijssen et al., 2018)). Many orthologs were unique to each gene set (Figure 2C); suggesting that our nervous-system wide profiling of the synaptic transcriptome has found new synaptically-localized transcripts in multiple neuron subtypes.

GO analysis of orthologs unique to the “Synaptic/Axonal Protein” and “Axon-Seq” list (Grey and Magenta Circles, Figure 2C) indicated that these genes were indeed synaptic in function (Figure 2D): previously-identified mammalian synaptic proteins are involved in *signaling receptor binding*, including a number of molecules involved in Wnt/Frizzled signaling (*dsh-2/Dvl1-3, egl-20/Wnt16)* and fibroblast growth receptor signaling (*egl-15/Fgfr1-4), cytoskeletal protein binding* including several kinesins *(klc-1/Klc2, klp-15;klp-16/Kifc3)*, and genes that bind anions and kinases. Axon-seq-unique orthologs also included structural and cytoskeletetal-regulating molecules (Figure 2D). Due to the relatively low number of orthologs (29, Figure 2C) that are unique to the synaptic neuropil list, we were unable to detect GO term enrichment; however, a number or these genes are receptor and membrane-associated (*yop-1/Reep6, Y57E12A.1/Serinc2, flap-1/Lrrfip2)*, microtubule associated (*C14H10.2/Jakmip1;Jakmip2*), and intracellular signaling molecules (*Y105E8A.2/Arhgef2, skr-2/Skp1*). The large number of genes involved in signaling and cytoskeletal regulation present in synaptic DEGS suggests that they are important for the dynamic remodeling that occurs in axons and synapses following unique stimuli.

### Synaptic DEGS are enriched for translational regulators and mRNA binding proteins

*C. elegans* synaptic DEGS orthologs shared between datasets may have an especially important role in regulating synaptic function. GO analysis of all genes shared between any two datasets (Axonal/synaptic protein, Axon-seq, and Synaptic Neuropil) revealed expected terms such as *actin binding, actin filament binding*, and *motor activity* (Figure 2D). The most significantly enriched GO term for the shared orthologs was *RNA binding*, with *mRNA 3’ UTR binding* and *translation regulator activity* also represented in the shared data set (Figure 2D). We determined which genes contributed to these GO terms and ranked them by synaptic enrichment (fold change relative to soma, Figure 3A). Translation regulators included five eukaryotic initiation factors (eIFs, *iff-1/eIF5A, ife-1/eIF4E, drr-2/eIF4H, ife-3/eIF4E, gcn-2/eIF2AK4*), two ribosomal subunits, and a eukaryotic elongation factor (*eef-1B.2/eEF1B2*). Multiple *C. elegans* orthologs of mammalian RNA binding proteins, including RNA Binding Motif protein 3/Cold inducible RNA binding protein (*rbm-3.1, rbm-3.2)* and Y-box binding protein 3 (*cey-2, cey-3*) were also present. Interestingly, the most significantly enriched and most numerous RNA binding proteins present in the synaptic DEGs were *pufs* (*puf-3, puf-5, puf-7, puf-8, puf-11;* Figure 3A), which are orthologs of mammalian *Pumilio1and Pumilio2 (Pum1/2).*

**Figure 3.**
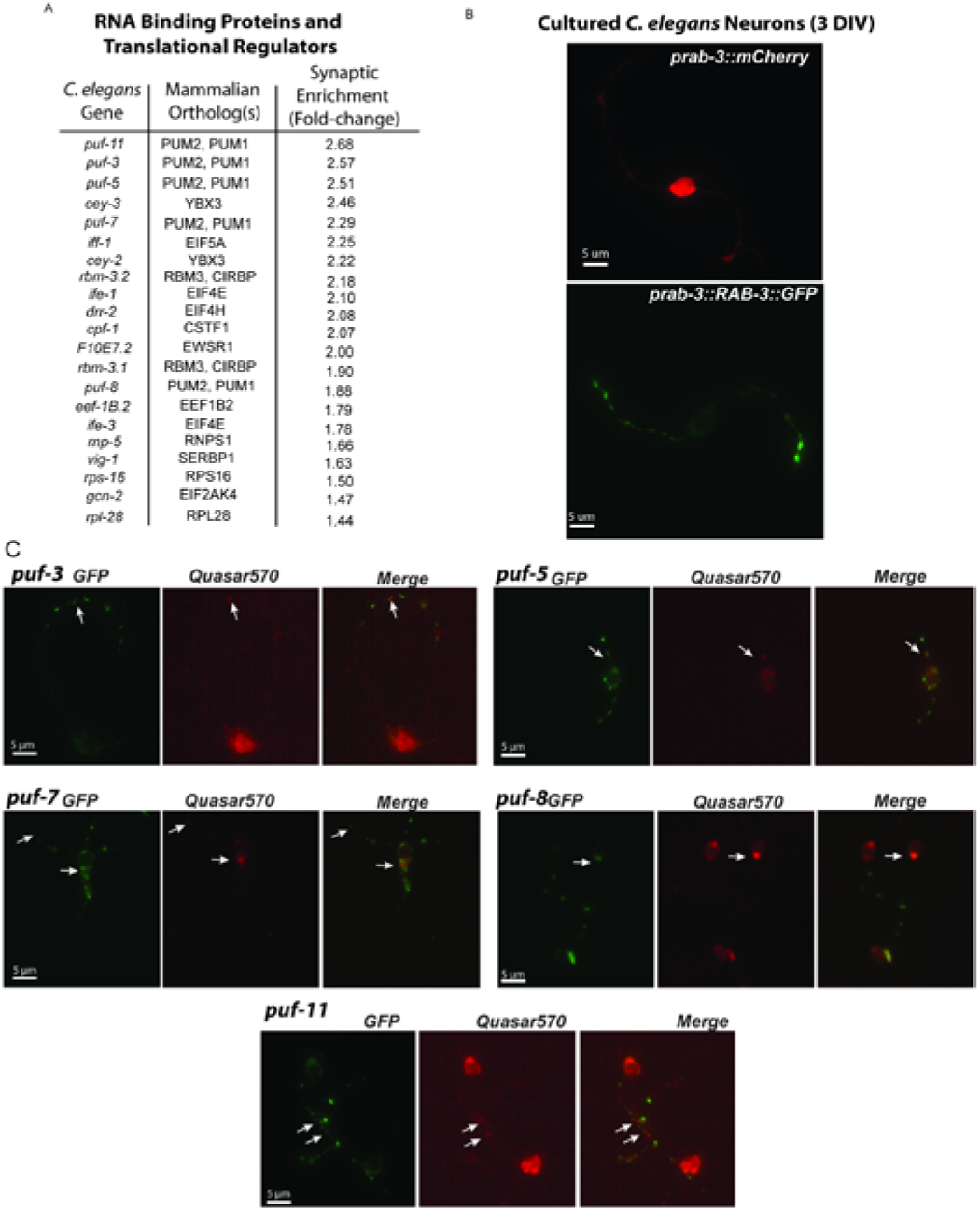
*puf* mRNAs are neuronally and axonally localized. **A)** List of translational regulators and RNA binding proteins present in synaptic DEGs, ranked by synaptic enrichment. **B)** Isolated *C. elegans* neurons 3 days *in vitro* (DIV), with somatic (*prab-3∷mCherry)* and synaptic (*prab-3∷RAB-3∷GFP)* markers. **C)** smFISH using Quasar570-labeled probes against individual *puf* mRNAs in isolated *prab-3∷RAB-3∷GFP* neurons.

### *C. elegans pumilio* mRNAs are expressed in neurons and axonally localized

Although mammalian PUM1/2 are expressed in neurons, *C. elegans* PUFs have primarily been characterized in the germline, where they regulate processes such as germ cell development, germline proliferation, and oocyte maturation (Wang et al., 2018). To confirm our sequencing data, we examined whether *puf* mRNAs were present in neurons and co-localized with the presynaptic marker RAB-3∷GFP. We cultured isolated *C. elegans* neurons from transgenic worms expressing *rab-3p∷mCherry* (soma label, Figure 3B) and *rab-3p∷RAB-3∷GFP* (synaptic label, Figure 3B). mCherry signal was primarily detected in the nucleus and soma of cultured neurons, while RAB-3∷GFP was detected in distinct puncta along neuronal projections (Figure 3B). To visualize mRNAs, we performed single molecule fluorescent *in situ* hybridization (smFISH (Raj et al., 2008)) on cultured *C. elegans* neurons using probes designed against individual *puf* mRNAs. We confirmed that all synaptically-enriched *puf* mRNAs identified by RNA-seq are indeed expressed in neurons (Figure 3C), and that mRNA puncta for individual *puf* mRNAs co-localize with synaptic RAB-3∷GFP. These results suggest that specific *puf* mRNAs are transported to pre-synaptic regions, presumably in preparation for translation.

### *C. elegans pufs* are both necessary memory components and memory repressors

What processes might axonal *pufs* regulate? We previously found that molecules that regulate pre-synaptic transport and transmission are important for learning and associative memory formation (Arey et al., 2018; Kaletsky et al., 2016; Lakhina et al., 2015; Li et al., 2016). Furthermore, *pumilio* is an important regulator of long-term memory in *Drosophila* (Dubnau et al., 2003), and mammalian PUM1/2 regulate dendrite morphogenesis, synaptic function, neuronal excitability, and hippocampal neurogenesis (Siemen et al., 2011; Vessey et al., 2010; Zhang et al., 2017). We therefore assessed the role of presynaptic *pufs* in positive olfactory associative memory, in which worms form a positive association with the neutral odorant butanone after pairing with food. A single food-butanone pairing results in a translation-dependent, intermediate-term memory one hour post-training that is forgotten in an active, translation-dependent manner by two hours post-training (Kauffman et al., 2010; Stein and Murphy, 2014). Adult-specific, RNAi-mediated knockdown of *puf-3* and *puf-5* in neuronal-RNAi-sensitive animals resulted in selective intermediate-term memory deficits (Figure 4A,B), while translation-independent learning and short-term memory were unaffected by knockdown (Figure S2A-D). These results suggested that *puf-3* and *puf-5* are essential memory promoting factors, which is in agreement with previous findings in *Drosophila* (Dubnau et al., 2003). However, adult-only RNAi-mediated knockdown of *puf-7* and *puf-8* resulted in enhanced memory: a positive association with butanone is maintained two hours post-training when compared to vector-control treated animals (Figure 4C,E). *puf-7* and *puf-8* deletion mutants replicated the extended memory phenotype of RNAi-treated animals, indicating that these two PUFs function as memory repressors.

**Figure 4.**
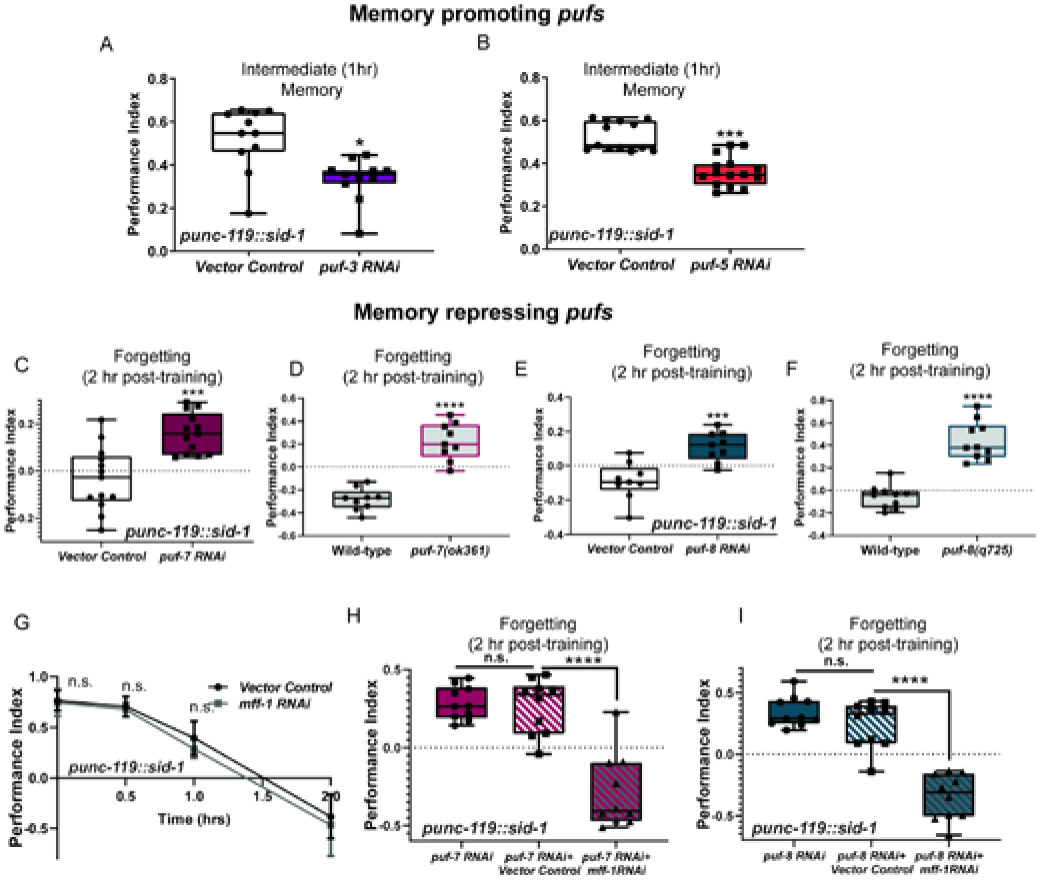
*puf* family members differentially regulate associative memory. **A-B)** *puf-3* and *puf-5* are required for normal intermediate-term memory formation. Mean ± SEM. n ≥ 8-10 per genotype. *p<0.05, ***p<0.001. **C-F)** RNAi-mediated knockdown or mutations in *puf-7* **(C-D)** and *puf-8* **(E-F)** result in an extended memory phenotype. **G)** *mff-1* RNAi treatment in neuronal RNAi-sensitive animals has no detectable effect on short/intermediate-term learning and memory. **H-I)** The mitochondrial fission factor *mff-1* is downstream of *puf-7* and *puf-8* in memory maintenance. Knockdown of *mff-1* and *puf-7* **(H)** and *puf-8* **(I)** suppresses the memory extending effects of *puf-7* **(H)** and *puf-8* **(I)** alone. Combining *puf-7* **(H)** and *puf-8* **(I)** RNAi with empty vector has no effect on behavior. Mean ± SEM. n ≥ 8-10 per genotype. *p<0.05, ***p<0.001, **** p<0.0001, n.s. p>0.05.

### Regulator of mitochondrial dynamics *mff-1* is downstream of the memory repressors PUF-7 and PUF-8

We wondered how PUF-7 and PUF-8 might act as inhibitors of memory formation. PUF-8, which is conserved in mammals, regulates mitochondrial dynamics and mitophagy by repressing translation of the mitochondrial fission factor *mff-1* (D’Amico et al., 2019), and reduction of PUF-8 function rescues age-related mitochondrial morphology in aged worms (D’Amico et al., 2019). Recently, axonal translation of mitochondrial proteins was found to be important for mitochondrial and axonal integrity in *Xenopus* retinal ganglion cells (Cioni et al., 2019). Because proper mitochondrial function is also necessary for synaptic transmission and plasticity (Alnaes and Rahamimoff, 1975), we hypothesized that PUF-7 and PUF-8 may inhibit memory formation by repression of *mff-1*. To test this hypothesis, we tested whether *mff-1* was necessary for the memory-promoting effects of *puf-7* and *puf-8* knockdown in adult animals. While RNAi-mediated knockdown of *mff-1* in adult, neuronal-RNAi-sensitive animals had no detectable effects on associative learning and memory when compared to control animals (Figure 4G), combining knockdown of *mff-1* and *puf-7* or *puf-8* suppressed the memory-promoting effects of either knockdown of *puf-7/8* or *puf-7/8* combined with empty vector control RNAi (Figure 4H,I). These results suggest that regulation of the inhibition of *mff-1* translation underlies the memory-repressing actions of *puf-7/8.*

## Discussion

Here we have defined the adult, nervous-system-wide presynaptic transcriptome for the first time in any system. We find that transcripts localized to synaptic regions reflect specialized neuronal functions that are expected in genes that would regulate plasticity. Synaptic DEGs, which are the most enriched in synaptic relative to somatic regions include receptors, molecular motors, and regulators of cytoskeletal remodeling and axon outgrowth, all of which are critical components of synaptic remodeling that occurs during plasticity.

We also found an unexpectedly large number of translational regulators and RNA-binding proteins in the list of synaptic DEGs. One possibility for this abundance is to enable stimulus-specific responses in neurons: regulating the expression of translational regulators can provide an additional layer of translational control. This is supported by the finding that following nerve injury, mTOR is rapidly translated to regulate the translation of other repair factors (Terenzio et al., 2018). *Puf* RNA binding proteins, orthologs of mammalian PUM1/2, are especially enriched in the synaptic DEG list. We confirmed that these *puf* mRNAs are indeed neuronal and co-localize with synaptic markers. Another *C. elegans pumilio* ortholog, *fbf-1*, has been previously shown to function in neurons and regulate behavior (Kaye et al., 2009; Stein and Murphy, 2014), but this is the first time that *puf* members of the *pumilio* gene family members have been found *C. elegans* neurons. It is not entirely unexpected that *pumilios* are in axonal regions. FMRP, which is known to interact with PUM2 (Zhang et al., 2017), is localized in axons during synaptogenesis (Parvin et al., 2019), so it is likely that other FMRP partners also exhibit axonal localization in higher organisms.

We find that synaptically-localized *pufs* differentially regulate associative memory: *puf-3/5* promote memory while *puf-7/8* are memory repressors. We determined that the mitochondrial fission factor *mff-1* is necessary for the memory repressing activity of *puf-7/8*, and by regulating its translation they can regulate mitochondrial dynamics. Translational regulation of mitochondrial proteins has already been shown to act in the axons of retinal ganglion cells, through RNA granule clustering on late endosomes (Cioni et al., 2019). Because mitochondria themselves power local translation and plasticity (Jang et al., 2016; Rangaraju et al., 2019), altering their dynamics through translational repression could negatively impact plasticity and memory. In the future it will be interesting to determine how *puf-3/5* and *puf-7/8* exert their differential effects on memory. It is known that despite their similarity, *pufs* in the *C. elegans* bind different targets and have different effects on oogenesis (Hubstenberger et al., 2012). It is likely that a similar mechanism underlies their differential memory effects.

By identifying the presynaptic transcriptome, we have demonstrated that presynaptic transcripts contribute to associative behaviors. Because many of these transcripts we identified have conserved functions in mammals, these findings set the framework for future studies for understanding the role that these presynaptic proteins play in plasticity, behavior and repair.

## Methods

### EXPERIMENTAL MODEL AND SUBJECT DETAILS

#### C. elegans genetics

All strains were maintained at 20°C on plates made from standard nematode growth medium (NGM: 3 g/L NaCl, 2.5 g/L Bacto-peptone, 17 g/L Bacto-agar in distilled water, with 1mL/L cholesterol (5 mg/mL in ethanol), 1 mL/L 1M CaCl_2_, 1 mL/L 1M MgSO_4_, and 25 mL/L 1M potassium phosphate buffer (pH 6.0) added to molten agar after autoclaving; (Brenner, 1974) or high growth medium (HGM: NGM recipe modified as follows: 20 g/L Bacto-peptone, 30 g/L Bacto-agar, and 4mL/L cholesterol (5 mg/mL in ethanol); all other components same as NGM), with OP50 E. coli as the food source. Experiments that did not involve RNAi treatments were performed using NGM and HGM plates seeded with OP50 E. coli for *ad libitum* feeding (Brenner, 1974); for RNAi experiments, the standard HGM molten agar was supplemented with 1?mL/L 1M IPTG (isopropyl β-d-1-thiogalactopyranoside) and 1mL/L 100mg/mL carbenicillin, and plates were seeded with HT115 E. coli for *ad libitum* feeding. Hypochlorite-synchronization to developmentally synchronize experimental animals was performed by collecting eggs from gravid hermaphrodites via exposure to an alkaline-bleach solution (*e.g.*, 8.0 mL water, 0.5 mL 5N KOH, 1.5 mL sodium hypochlorite), followed by repeated washing of collected eggs in M9 buffer (6 g/L Na_2_HPO_4_, 3 g/L KH_2_PO_4_, 5 g/L NaCl and 1 mL/L 1M MgSO_4_ in distilled water; (Brenner, 1974)). For RNAi experiments, animals were transferred at the L4 larval stage onto HGM-RNAi plates until Day 2 of adulthood, when the animals were subjected to behavioral testing.

##### Strains

Wild-type: (N2 Bristol); Transgenic strains: NM2415 (*lin-15B(n765); jsIs68[Prab-3∷GFP∷rab-3 + lin-15(+)]*), LC108 (*vIs69 [pCFJ90(Pmyo-2∷mCherry + Punc-119∷sid-1)]*) OH441(*punc-119∷GFP)*; Mutant strains: RB652 (*puf-7(ok361*)) and JK3231(*puf-8(q725))* were obtained from the *Caenorhabditis* Genetics Center (University of Minnesota, Minneapolis, MN). The transgenic strain CQ574 (*lin-15B(n765); jsIs682 [Prab-3∷GFP∷rab-3 + lin-15(+)]; wqIs3 [Prab3∷mCherry*]) was generated by UV integration (Mariol et al., 2013) of NM2415 (*lin-15B(n765); jsIs68[Prab-3∷GFP∷rab-3 + lin-15(+)]*) animals microinjected with a *Prab3∷mCherry* transgenic construct, followed by 3 rounds of outcrossing with wild-type (N2 Bristol) worms.

### METHOD DETAILS

#### Adult cell isolation

Adult cell isolation was performed as described previously (Kaletsky et al., 2016). Synchronized day 1 adult CQ574 (*lin-15B(n765); jsIs682 [Prab-3∷GFP∷rab-3 + lin-15(+)]; wqIs3 [Prab3∷mCherry*]) worms washed with M9 buffer to remove excess bacteria. The pellet (∼ 250 µl) was washed with 500 µl lysis buffer (200 mM DTT, 0.25% SDS, 20 mM Hepes pH 8.0, 3% sucrose) and resuspended in 1000 µl lysis buffer. Worms were incubated in lysis buffer with gentle rocking for 6.5 minutes at room temperature. The pellet was washed 6× with M9 and resuspended in 20 mg/ml pronase from *Streptomyces griseus* (Sigma-Aldrich). Worms were incubated at room temperature (<20 minutes) with periodic mechanical disruption by pipetting every 2 min. When most worm bodies were dissociated, leaving only small debris and eggs, ice-cold PBS buffer containing 2% fetal bovine serum (Gibco) was added. RNA from FACS-sorted neurons was prepared for RNA-seq and subsequent analysis (see FACS isolation and RNA seq Analysis for more details).

#### FACS isolation of dissociated cells

Cells were briefly subjected to SDS-DTT treatment, proteolysis, mechanical disruption, cell filtering, FACS, RNA amplification, library preparation, and single-end (180 nt) Illumina sequencing, as previously described. Neuron cell suspensions were passed over a 5 μm syringe filter (Millipore). The filtered cells were diluted in Osmo-balance Leibovitz L-15/2% FBS and sorted using a FACSVantage SE w/ DiVa (BD Biosciences; 488nm excitation for GFP detection, 568nm excitation for mCherry detection). Gates for detection were set by comparison to non-fluorescent N2 cell suspensions prepared on the same day from a population of worms synchronized alongside the experimental samples. Positive fluorescent events were sorted directly into Eppendorf tubes containing Trizol LS for subsequent RNA extraction. For each sample, approximately 30,000–130,000 GFP or mCherry positive events were collected, yielding 5–25 ng total RNA.

#### RNA isolation, amplification, library preparation, and sequencing

RNA was isolated from FACS-sorted samples as previously described (Kaletsky et al., 2016; Kaletsky et al., 2018). Briefly, RNA was extracted using standard Trizol/ chloroform/ isopropanol method, DNase digested, and cleaned using Qiagen RNEasy Minelute columns. Agilent Bioanalyzer RNA Pico chips were used to assess quality and quantity of isolated RNA. RNA sequencing libraries were prepared directly from quality assessed RNA using the SMARTer Stranded Total RNA kit v2-Pico input mammalian, as per manufacturer suggested practices. The resultant sequencing libraries were then submitted for sequencing on the Illumina HiSeq 2000 platform. ∼ 75–190 million reads (average of 128,910,533 reads) were obtained for each sample and mapped to the *C*. *elegans* genome. Raw sequencing reads are available at NCBI Bioproject: PRJNA559377.

#### Microscopy

Imaging of day 1 CQ574 (*lin-15B(n765); jsIs682 [Prab-3∷GFP∷rab-3 + lin-15(+)]; wqIs3 [Prab3∷mCherry*]) adults was performed on a Nikon A1 confocal microscope at 60× magnification, and *z* stacks were processed in Nikon NIS elements software. For imaging of smFISH samples, Z-stack multi-channel (DIC, TRITC, GFP) of isolated neurons were imaged every 0.2 µm at 100X magnification on a Nikon Eclipse Ti inverted microscope; Maximum Intensity Projections and 3D reconstructions of neurons were built with Nikon *NIS-Elements*.

#### C. elegans neuronal cell isolation and culture

Isolation and culture of neurons was performed as previously described (Zhang et al., 2003), with modifications. Animals were synchronized by hypochlorite treatment and grown on OP50-seeded NGM or HGM plates until the L4 larval stage. One hour prior to cell isolation, animals were allowed to incubate in M9 buffer to clear the gut of bacteria. After one hour, animals were incubated in 200-300 µl freshly thawed sterile SDS-DTT solution (200 mM DTT, 0.25% SDS, 20 mM HEPES, pH 8.0, 3% sucrose, stored at −20°C) for 4 min at room temperature. The animals were washed 3-5 times in M9 buffer and pelleted by centrifugation in a tabletop centrifuge. Animals were then digested in 15 mg/ml pronase for 20–25 min and subjected to mechanical disruption by frequent pipetting. Pronase digestion was stopped by adding 900 µl L-15 medium (Invitrogen, Carlsbad, CA) supplemented with 10% fetal bovine serum (Invitrogen, Carlsbad, CA), 50 U/ml penicillin, and 50 µg/ml streptomycin (Sigma-Aldrich, St. Louis, MO) and adjusted to 340 mOsm. Cells were pelleted by centrifugation at 10,000 rpm for 5 min at 4°C, and washed 2 times with L-15/FBS. The pellet was resuspended with 1 ml L-15/FBS and settled on ice for 30 min. The top 800 µl cell suspension devoid of large worm debris was transferred to a new tube and pelleted by centrifugation at 10,000 rpm for 5 min at 4°C. Cell pellets were resuspended in fresh L-15/FBS. 40 to 50 µl of cell suspension was plated onto the center an acid-washed coverslip coated with 0.5 mg/ml peanut lectin (Sigma-Aldrich, St. Louis, MO). Cells were allowed to adhere overnight in a 20°C incubator without CO_2_ in Snapware plastic containers and humidified with moist paper towels. 24 hours later, debris and unbound cells were washed off with L-15, and 1ml of fresh L-15/FBS was added to the coverslips. After 3 days in culture, smFISH was performed.

#### Single molecule fluorescence in situ hybridization

Custom Stellaris® FISH Probes were designed against *puf-3, puf-5, puf-7, puf-8*, and *puf-11* by utilizing the Stellaris® RNA FISH Probe Designer (Biosearch Technologies, Inc., Petaluma, CA) available online at www.biosearchtech.com/stellarisdesigner (version 4.2). The isolated neurons from *C. elegans* strain NM2415 (*lin-15B(n765); jsIs68[Prab-3∷GFP∷rab-3 + lin-15(+)]*) were hybridized with individual *puf* Stellaris RNA FISH Probe sets labeled with Quasar 570 (Biosearch Technologies, Inc.), following the manufacturer’s instructions available online at www.biosearchtech.com/stellarisprotocols. Briefly, cells were fixed with a paraformaldehyde fixation buffer (3.7% paraformaldehyde in 1X PBS) for 10 minutes at room temperature, washed twice with 1X PBS, and permeabilized for at least 1 hour with 70% EtOH. After permeabilization, cells were washed with Wash Buffer A (Biosearch Technologies, Inc.) containing 10% deionized formamide (Sigma Aldrich), and hybridized overnight at 37°C with Stellaris RNA FISH Probe sets at a final concentration of 250nM in Hybridization buffer (Biosearch Technologies, Inc.) containing 10% deionized formamide (Sigma Aldrich). After hybridization, cells are washed twice at 37°C Wash Buffer A (Biosearch Technologies, Inc.) containing 10% deionized formamide (Sigma Aldrich), followed by a 2-5 minute wash at room temperature with Wash Buffer B (Biosearch Technologies, Inc.). Coverslips were then applied to slides and imaged (see *Microscopy* section).

#### Behavioral Assays

Wild-type, mutant, and transgenic animals were trained and tested for or short/intermediate term memory as previously described (Kauffman et al., 2010). Briefly, synchronized day 1 adult hermaphrodites (or day 2 adult worms for RNAi treated animals) were washed from HGM plates with M9 buffer, allowed to settle by gravity, and washed again with M9 buffer. After washing, the animals are starved for 1 hr in M9 buffer. For 1 CS-US pairing, worms were then transferred to 10 cm NGM conditioning plates (seeded with OP50 *E. coli* bacteria and with 6μl 10% 2-butanone (Acros Organics) in ethanol on the lid) for 1 hr. After conditioning, the trained population of worms were tested for chemotaxis to 10% butanone vs. an ethanol control either immediately (0 hr) or after being transferred to 10 cm NGM plates with fresh OP50 for specified intervals before testing (30 mins-2 hrs), using standard, previously described chemotaxis assay conditions (Bargmann et al., 1993).

Chemotaxis indices were calculated as follows: **(#worms**_**Butanone**_ **-#worm**_**Ethanol**_**)/(Total #worms)**. Performance index is the change in chemotaxis index following training relative to the naïve chemotaxis index. The calculation for Performance Index is: **Chemotaxis Index**_**Trained**_ **-Chemotaxis Index**_**Naive**_.

### QUANTIFICATION AND STATISTICAL ANALYSIS

#### RNA-seq data analysis

FASTQC was used to inspect the quality scores of the raw sequence data, and to look for biases. Reads were mapped to the *C*. *elegans* genome (WormBase 245) using STAR with WormBaseID gene model annotations (using default parameters). Count matrices were generated for the number of reads overlapping with the gene body of protein coding genes using htseqCounts. DESeq2 was used for differential expression analysis and the principal components analysis. Genes at FDR = 0.05 were considered significantly differentially expressed.

#### Gene ontology analysis

Hypergeometric tests of Gene Ontology terms were performed on tissue-enriched gene lists using g:Profiler; GO terms reported are a significance of *q-value* < 0.05 unless otherwise noted.

#### Behavioral Assay Analysis

For the comparison of performance indices between two genotypes (i.e. *puf-7(ok361)* vs wild-type) or two RNAi treatments (i.e. Vector control RNAi and *puf-3* RNAi), two-tailed unpaired Student’s t-tests with Welch’s corrections were used. When 3 or more groups of RNAis were compared, one-way analysis of variances followed by Bonferroni post hoc tests for multiple comparisons were performed. 2-way ANOVAs were used for evaluating effects between RNAi treatment (*mff-1* and Vector Control) and timepoint (0hr, 0.5hr, 1hr, 2hr) on performance indices with a significant interaction between factors (*p* < 0.0001) leading to the performance of Bonferroni post-hoc comparisons to determine differences between individual groups. Experiments were repeated on separate days with separate populations, to confirm that results were reproducible. Prism 8 software was used for all statistical analyses. Additional statistical details of experiments, including sample size (with n representing the number of chemotaxis assays performed for behavior, and number of cells imaged for microscopy), can be found in the figure legends.

## Acknowledgments

We thank the *C. elegans* Genetics Center for strains; the Genomics Core Facility at Princeton University; Christina DeCoste, Katherine Rittenbach and the Flow Cytometry Core at Princeton University; and Lance Parsons for assistance in generating graphical representation of Synaptic DEGs. CTM is the Director of the Glenn Center for Aging Research at Princeton and an HHMI-Simons Faculty Scholar.

## Funding

Support for this work was provided by a DP1 Award (NIGMS 1DP2OD004402-01) and NIA R01 (NIA 5R01AG034446) to CTM, The Glenn Foundation for Medical Research (GMFR CNV1001899), and the HHMI Faculty Scholar Program (AWD1005048), and an F32 NRSA (NIA 5F32AG046106) to RNA.

## Author contributions

RNA, RK, and CTM designed experiments. RK constructed the dual-labelled transgenic line. RNA and RK performed experiments and analyzed data. RNA, RK, and CTM wrote the manuscript.

### Competing interests

Authors declare no competing interests.

---

**Supplementary Figure S1.**
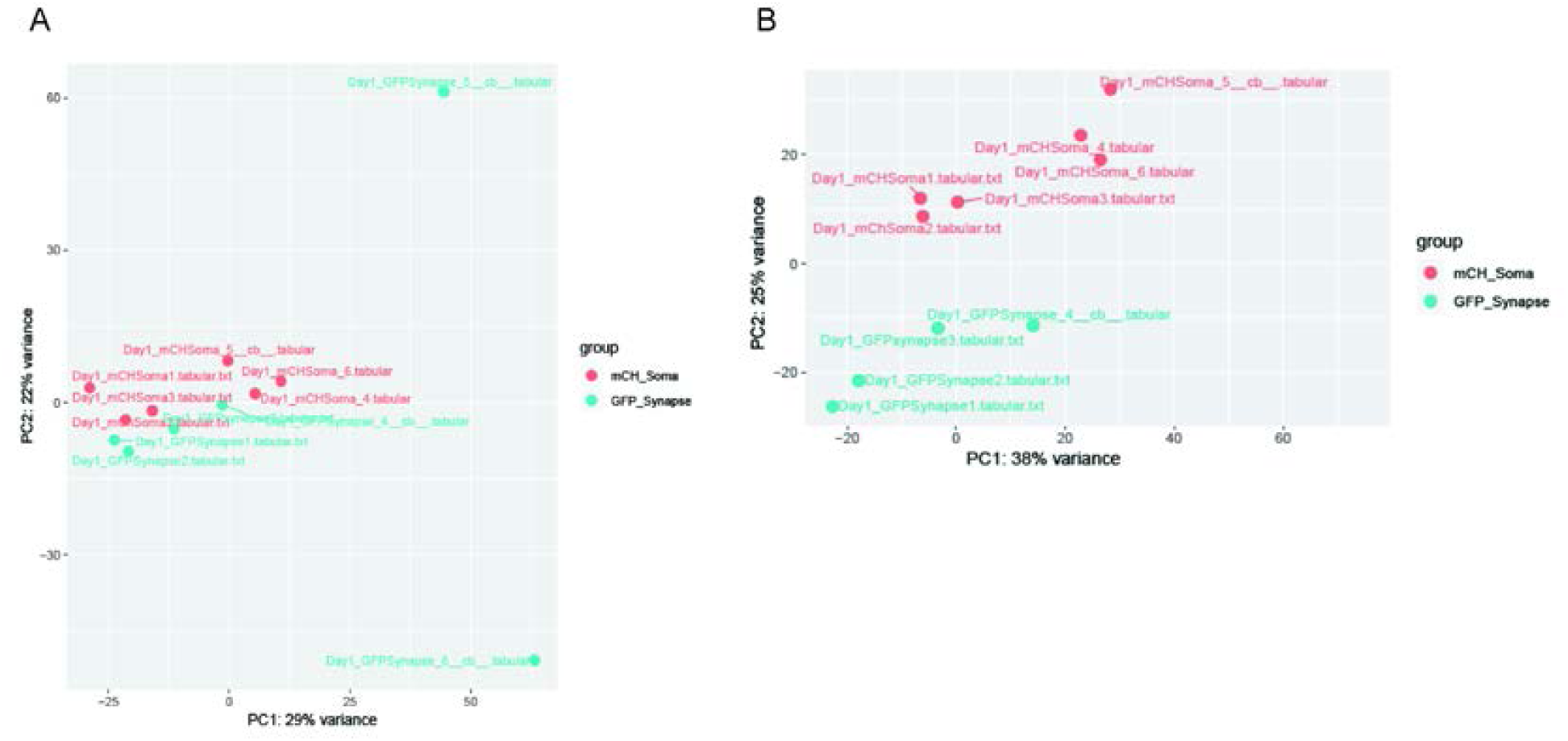
**A)** Principle components analysis of six mCherry+ (red) somatic and GFP+ (blue) synaptic RNA-seq samples performed as part of DESeq2 analysis. **B)** Principle component analysis of four synaptic (blue) and six somatic (red) RNA-seq samples after removal of outliers. Remaining samples cluster by subcompartment.

**Supplementary Figure S2.**
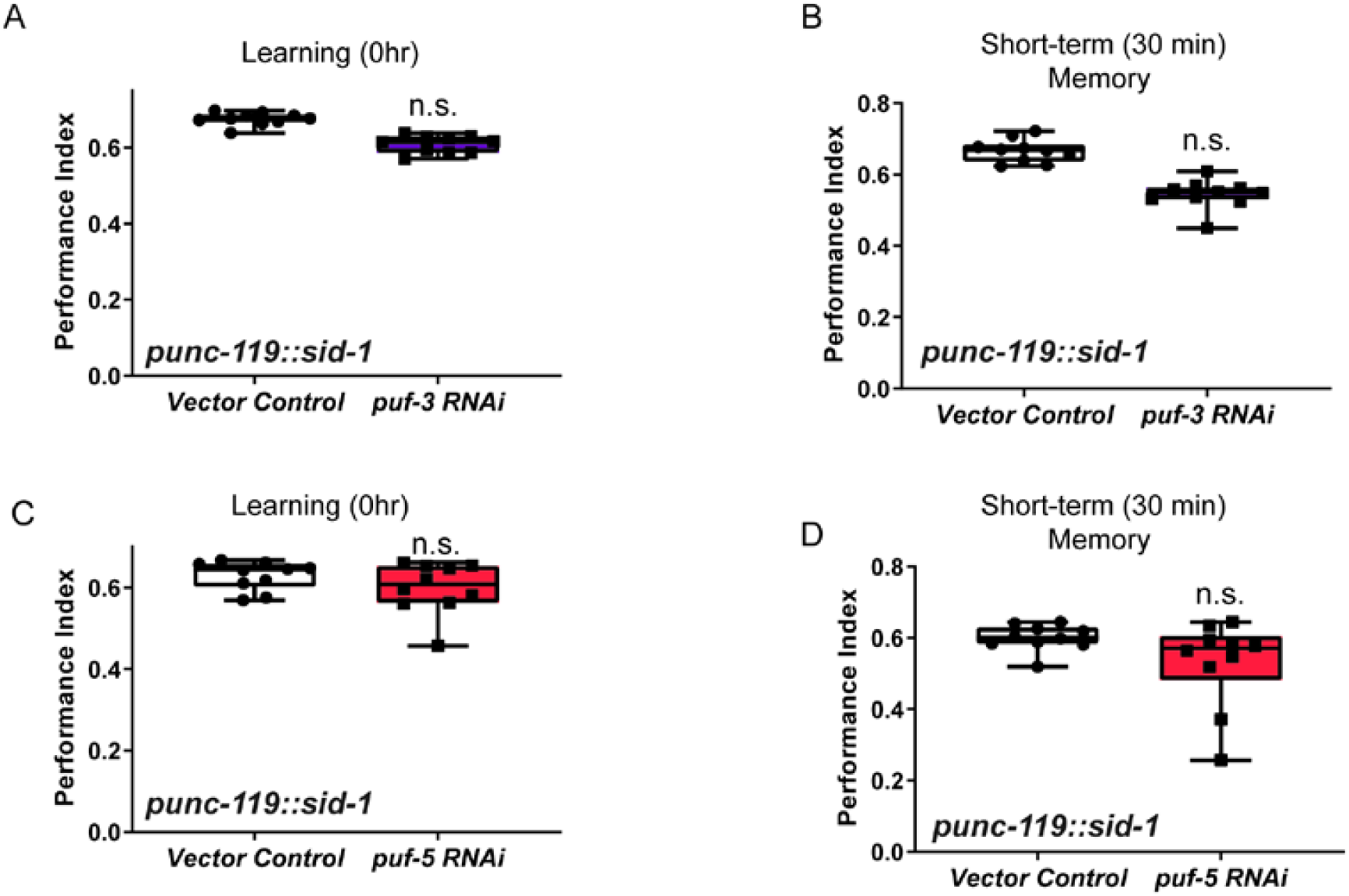
**A-D)** *puf-3 (***A,B**) and *puf-5* (**C,D**) RNA treatment in neuronal RNAi-sensitive animals has no detectable effect on learning (**A,C**) or short-term memory (**B,D**).

**Supplemental Table 1. Synapse-expressed genes identified by RNA-seq.**

**Supplemental Table 2. DESeq2 Results comparing GFP+ Synaptic samples to mCherry+ Somatic samples from FACS sorted dual-fluorescent lines.**

**Supplemental Table 3. Mammalian orthologs of synaptic DEGs that have been previously identified to function in synaptic and axonal regions, were identified by axon-seq (Njissen et al., 2012), or by analysis of the hippocampal synaptic neuropil (Cajigas et al., 2012).**

## References

Alnaes, E., and Rahamimoff, R. (1975). On the role of mitochondria in transmitter release from motor nerve terminals. J Physiol 248, 285–306.

Arey, R.N., Stein, G.M., Kaletsky, R., Kauffman, A., and Murphy, C.T. (2018). Activation of G. Neuron 98, 562–574.e565.

Bargmann, C.I., Hartwieg, E., and Horvitz, H.R. (1993). Odorant-selective genes and neurons mediate olfaction in C. elegans. Cell 74, 515–527.

Batista, A.F.R., Martínez, J.C., and Hengst, U. (2017). Intra-axonal Synthesis of SNAP25 Is Required for the Formation of Presynaptic Terminals. Cell Rep 20, 3085–3098.

Brenner, S. (1974). The genetics of Caenorhabditis elegans. Genetics 77, 71–94.

Cajigas, I.J., Tushev, G., Will, T.J., tom Dieck, S., Fuerst, N., and Schuman, E.M. (2012). The local transcriptome in the synaptic neuropil revealed by deep sequencing and high-resolution imaging. Neuron 74, 453–466.

Cioni, J.M., Lin, J.Q., Holtermann, A.V., Koppers, M., Jakobs, M.A.H., Azizi, A., Turner-Bridger, B., Shigeoka, T., Franze, K., Harris, W.A., and Holt, C.E. (2019). Late Endosomes Act as mRNA Translation Platforms and Sustain Mitochondria in Axons. Cell 176, 56–72.e15.

Cracco, J.B., Serrano, P., Moskowitz, S.I., Bergold, P.J., and Sacktor, T.C. (2005). Protein synthesis-dependent LTP in isolated dendrites of CA1 pyramidal cells. Hippocampus 15, 551–556.

D’Amico, D., Mottis, A., Potenza, F., Sorrentino, V., Li, H., Romani, M., Lemos, V., Schoonjans, K., Zamboni, N., Knott, G., et al. (2019). The RNA-Binding Protein PUM2 Impairs Mitochondrial Dynamics and Mitophagy During Aging. Mol Cell 73, 775–787.e710.

Dubnau, J., Chiang, A.S., Grady, L., Barditch, J., Gossweiler, S., McNeil, J., Smith, P., Buldoc, F., Scott, R., Certa, U., et al. (2003). The staufen/pumilio pathway is involved in Drosophila long-term memory. Curr Biol 13, 286–296.

Glock, C., Heumüller, M., and Schuman, E.M. (2017). mRNA transport & local translation in neurons. Curr Opin Neurobiol 45, 169–177.

Hafner, A.S., Donlin-Asp, P.G., Leitch, B., Herzog, E., and Schuman, E.M. (2019). Local protein synthesis is a ubiquitous feature of neuronal pre- and postsynaptic compartments. Science 364.

Huber, K.M., Kayser, M.S., and Bear, M.F. (2000). Role for rapid dendritic protein synthesis in hippocampal mGluR-dependent long-term depression. Science 288, 1254–1257.

Hubstenberger, A., Cameron, C., Shtofman, R., Gutman, S., and Evans, T.C. (2012). A network of PUF proteins and Ras signaling promote mRNA repression and oogenesis in C. elegans. Dev Biol 366, 218–231.

Jang, S., Nelson, J.C., Bend, E.G., Rodríguez-Laureano, L., Tueros, F.G., Cartagenova, L., Underwood, K., Jorgensen, E.M., and Colón-Ramos, D.A. (2016). Glycolytic Enzymes Localize to Synapses under Energy Stress to Support Synaptic Function. Neuron 90, 278–291.

Jung, H., Yoon, B.C., and Holt, C.E. (2012). Axonal mRNA localization and local protein synthesis in nervous system assembly, maintenance and repair. Nat Rev Neurosci 13, 308–324.

Kaletsky, R., Lakhina, V., Arey, R., Williams, A., Landis, J., Ashraf, J., and Murphy, C.T. (2016). The C. elegans adult neuronal IIS/FOXO transcriptome reveals adult phenotype regulators. Nature 529, 92–96.

Kaletsky, R., Yao, V., Williams, A., Runnels, A.M., Tadych, A., Zhou, S., Troyanskaya, O.G., and Murphy, C.T. (2018). Transcriptome analysis of adult Caenorhabditis elegans cells reveals tissue-specific gene and isoform expression. PLoS Genet 14, e1007559.

Kang, H., and Schuman, E.M. (1996). A requirement for local protein synthesis in neurotrophin-induced hippocampal synaptic plasticity. Science 273, 1402–1406.

Kauffman, A.L., Ashraf, J.M., Corces-Zimmerman, M.R., Landis, J.N., and Murphy, C.T. (2010). Insulin signaling and dietary restriction differentially influence the decline of learning and memory with age. PLoS Biol 8, e1000372.

Kaye, J.A., Rose, N.C., Goldsworthy, B., Goga, A., and L’Etoile, N.D. (2009). A 3’UTR pumilio-binding element directs translational activation in olfactory sensory neurons. Neuron 61, 57–70.

Kim, W., Underwood, R.S., Greenwald, I., and Shaye, D.D. (2018). OrthoList 2: A New Comparative Genomic Analysis of Human and. Genetics 210, 445–461.

Lakhina, V., Arey, R.N., Kaletsky, R., Kauffman, A., Stein, G., Keyes, W., Xu, D., and Murphy, C.T. (2015). Genome-wide functional analysis of CREB/long-term memory-dependent transcription reveals distinct basal and memory gene expression programs. Neuron 85, 330–345.

Li, L.B., Lei, H., Arey, R.N., Li, P., Liu, J., Murphy, C.T., Xu, X.Z., and Shen, K. (2016). The Neuronal Kinesin UNC-104/KIF1A Is a Key Regulator of Synaptic Aging and Insulin Signaling-Regulated Memory. Curr Biol 26, 605–615.

Love, M.I., Huber, W., and Anders, S. (2014). Moderated estimation of fold change and dispersion for RNA-seq data with DESeq2. Genome Biol 15, 550.

Mahoney, T.R., Liu, Q., Itoh, T., Luo, S., Hadwiger, G., Vincent, R., Wang, Z.W., Fukuda, M., and Nonet, M.L. (2006). Regulation of synaptic transmission by RAB-3 and RAB-27 in Caenorhabditis elegans. Mol Biol Cell 17, 2617–2625.

Mariol, M.C., Walter, L., Bellemin, S., and Gieseler, K. (2013). A rapid protocol for integrating extrachromosomal arrays with high transmission rate into the C. elegans genome. J Vis Exp, e50773.

Martin, K.C., Casadio, A., Zhu, H., Yaping, E., Rose, J.C., Chen, M., Bailey, C.H., and Kandel, E.R. (1997). Synapse-specific, long-term facilitation of aplysia sensory to motor synapses: a function for local protein synthesis in memory storage. Cell 91, 927–938.

Nijssen, J., Aguila, J., Hoogstraaten, R., Kee, N., and Hedlund, E. (2018). Axon-Seq Decodes the Motor Axon Transcriptome and Its Modulation in Response to ALS. Stem Cell Reports 11, 1565–1578.

Parvin, S., Takeda, R., Sugiura, Y., Neyazaki, M., Nogi, T., and Sasaki, Y. (2019). Fragile X mental retardation protein regulates accumulation of the active zone protein Munc18-1 in presynapses via local translation in axons during synaptogenesis. Neurosci Res 146, 36–47.

Raj, A., van den Bogaard, P., Rifkin, S.A., van Oudenaarden, A., and Tyagi, S. (2008). Imaging individual mRNA molecules using multiple singly labeled probes. Nat Methods 5, 877–879.

Rangaraju, V., Lauterbach, M., and Schuman, E.M. (2019). Spatially Stable Mitochondrial Compartments Fuel Local Translation during Plasticity. Cell 176, 73–84.e15.

Scarnati, M.S., Kataria, R., Biswas, M., and Paradiso, K.G. (2018). Active presynaptic ribosomes in the mammalian brain, and altered transmitter release after protein synthesis inhibition. Elife 7.

Shigeoka, T., Jung, H., Jung, J., Turner-Bridger, B., Ohk, J., Lin, J.Q., Amieux, P.S., and Holt, C.E. (2016). Dynamic Axonal Translation in Developing and Mature Visual Circuits. Cell 166, 181–192.

Siemen, H., Colas, D., Heller, H.C., Brüstle, O., and Pera, R.A. (2011). Pumilio-2 function in the mouse nervous system. PLoS One 6, e25932.

Stein, G.M., and Murphy, C.T. (2014). C. elegans positive olfactory associative memory is a molecularly conserved behavioral paradigm. Neurobiol Learn Mem 115, 86–94.

Terenzio, M., Koley, S., Samra, N., Rishal, I., Zhao, Q., Sahoo, P.K., Urisman, A., Marvaldi, L., Oses-Prieto, J.A., Forester, C., et al. (2018). Locally translated mTOR controls axonal local translation in nerve injury. Science 359, 1416–1421.

Tushev, G., Glock, C., Heumüller, M., Biever, A., Jovanovic, M., and Schuman, E.M. (2018). Alternative 3’ UTRs Modify the Localization, Regulatory Potential, Stability, and Plasticity of mRNAs in Neuronal Compartments. Neuron 98, 495–511.e496.

Twiss, J.L., and Fainzilber, M. (2009). Ribosomes in axons--scrounging from the neighbors? Trends Cell Biol 19, 236–243.

Vessey, J.P., Schoderboeck, L., Gingl, E., Luzi, E., Riefler, J., Di Leva, F., Karra, D., Thomas, S., Kiebler, M.A., and Macchi, P. (2010). Mammalian Pumilio 2 regulates dendrite morphogenesis and synaptic function. Proc Natl Acad Sci U S A 107, 3222–3227.

Wang, D.O., Kim, S.M., Zhao, Y., Hwang, H., Miura, S.K., Sossin, W.S., and Martin, K.C. (2009). Synapse- and stimulus-specific local translation during long-term neuronal plasticity. Science 324, 1536–1540.

Wang, M., Ogé, L., Perez-Garcia, M.D., Hamama, L., and Sakr, S. (2018). The PUF Protein Family: Overview on PUF RNA Targets, Biological Functions, and Post Transcriptional Regulation. Int J Mol Sci 19.

Younts, T.J., Monday, H.R., Dudok, B., Klein, M.E., Jordan, B.A., Katona, I., and Castillo, P.E. (2016). Presynaptic Protein Synthesis Is Required for Long-Term Plasticity of GABA Release. Neuron 92, 479–492.

Zhang, F., Phiel, C.J., Spece, L., Gurvich, N., and Klein, P.S. (2003). Inhibitory phosphorylation of glycogen synthase kinase-3 (GSK-3) in response to lithium. Evidence for autoregulation of GSK-3. J Biol Chem 278, 33067–33077.

Zhang, M., Chen, D., Xia, J., Han, W., Cui, X., Neuenkirchen, N., Hermes, G., Sestan, N., and Lin, H. (2017). Posttranscriptional regulation of mouse neurogenesis by Pumilio proteins. Genes Dev 31, 1354–1369.

